# Associations between brain structure and both language proficiency and language balance in early bilinguals

**DOI:** 10.64898/2026.05.14.725184

**Authors:** Mia R. Coutinho, Edith Brignoni-Pérez, Nasheed I. Jamal, Guinevere F. Eden

**Author notes:** **Corresponding author** Guinevere Eden, D.Phil., Professor, Department of Pediatrics Georgetown University Medical Center Suite 150, Building D 4000 Reservoir Road, NW Washington, DC 20057, USA.

## Abstract

Prior studies in bilinguals have reported relationships between brain structure and the dimensions of (i) language proficiency or (ii) language balance (the discrepancy between a bilingual’s two proficiencies), but rarely both, even though they are highly related. These studies were often conducted in late bilinguals and the analyses limited to regions of interest. Here, we tested for relationships between brain structure and these two dimensions in 46 early cultural Spanish-English bilinguals (mean age = 16.7 years) at the level of the whole brain for gray matter volume (GMV) and cortical thickness (CT). Results revealed a positive association between GMV and proficiency in the weaker language in the right angular gyrus (AG; BA 39) extending into the superior temporal gyrus (BA 22). More balanced bilingualism was also associated with more GMV in the AG (BA 39), in addition to less GMV in left postcentral gyrus (BA 1), right cerebellum lobule IX and right superior occipital gyrus (BA 18). However, these relationships between GMV and balance disappeared after controlling for language proficiency. No significant associations were observed for CT and these two dimensions of language. Our findings suggest that relationships between GMV and balance are driven by language proficiency, and that the relationship between GMV and language proficiency likely does not involve language-specific mechanisms, given the location of the association is in the right inferior parietal cortex. Together, this study separates the neuroanatomical bases of these two language dimensions and places them in brain regions outside those usually targeted in prior studies.

**Highlights:**

1. Neuroanatomy was correlated with proficiencies in early Spanish-English bilinguals
2. Right angular gyrus gray matter volume (GMV) was positively related to proficiency
3. GMV was positively related to balance, but not after controlling for proficiency
4. Relations with these language dimensions are located outside of language cortex
5. No significant associations were observed for cortical thickness

## 1 Introduction

The current study examined the relationship between brain structure and two interrelated dimensions that vary among bilinguals: proficiency in each language, and balance between the two languages. Language proficiency refers to the level of skill or competence and has been assessed in a variety of ways, (Claussenius-Kalman et al., 2021; Grogan et al., 2012; Mechelli et al., 2004; Nguyen et al., 2023; Xu et al., 2024) usually using objective performance measures to gauge vocabulary, naming, or comprehension, or a combination of such types of verbal skills (for simplicity, we here refer to any of these as a measure of language proficiency). A “balanced bilingual” refers to an individual whose level of proficiency for their two languages is similar or the same, whereas an “unbalanced bilingual” has a discrepancy in the proficiency of their languages. While language proficiency and balance are interrelated, across the spectrum of language proficiency bilinguals can show a range in their language balance scores, such that even a bilingual who is considered proficient in both languages can nevertheless have a sizable gap between these, rendering them unbalanced.

Prior studies in bilinguals have reported on relationships between brain structure and language proficiency (Claussenius-Kalman et al., 2021; Grogan et al., 2012; Mechelli et al., 2004; Nguyen et al., 2023; Xu et al., 2024). A positive relationship has usually been attributed to the language learning experience (Grogan et al., 2012; Mechelli et al., 2004). Others have studied the relationships between brain structure and language balance (Archila-Suerte et al., 2018; Claussenius-Kalman et al., 2021; Marin-Marin et al., 2022). Because these investigations have mostly been conducted separately, it is unknown whether relationships between brain structure and language proficiency are intertwined with relationships between brain structure and language balance. The current study builds on this work by directly disambiguating these relationships. Further, almost all of the work noted above was based on groups dominated by bilinguals who acquired their second language after age seven years, often in late childhood and some even in adolescence or adulthood. In such late bilinguals, bilingual status is often the result of intense, sustained direct education, and second language acquisition may be facilitated in those with an aptitude for language learning. In contrast, cultural bilinguals learn their languages early as part of their family environment, often from birth. Advancing our understanding on the nature of the relationship between brain structure and language proficiency as well as language balance in early cultural bilinguals, who are the focus of the current study, will inform the brain-based mechanisms that lead to language-learning.

Here, we consider prior investigations into relationships between brain structure and language proficiency in more detail. The earliest contribution was made by Mechelli and colleagues (2004), who found no correlations between gray matter density (GMD) and second language proficiency (a combination of verbal measures including reading, writing, and speech comprehension and production) at the level of the whole brain, but found a positive correlation in a region of interest (ROI) analysis of the left inferior parietal cortex (IPC) in a sample of mostly late bilinguals. Using a whole-brain analysis, Grogan et al. (2012) found GMD and second-language proficiency (measured by lexical efficiency) to be positively correlated in the left inferior frontal cortex in bilinguals who acquired their languages during early or late childhood. However, ROI analyses in left and right supramarginal gyri and left posterior temporal cortex yielded no findings. When analyses were conducted for gray matter volume (GMV), there were no results (Grogan et al., 2012). In these studies, as is common across the literature, language abilities were based on the second language (in these cases, English) which in late bilinguals tends to be weaker than the first language. However, more recent studies have studied language performance measures for both of the bilingual’s languages (Nguyen et al., 2023; Xu et al., 2024). Xu and colleagues (2024) examined language proficiency (receptive picture vocabulary combined with either reading or listening comprehension and/or the expressive picture vocabulary) in both languages in a group that combined early and late bilinguals using an ROI approach within subcortical regions. They reported more GMV in the left accumbens, right thalamus and bilateral globus pallidus was related to greater language proficiency in the stronger language, while less GMV in the right accumbens and bilateral globus pallidus related to greater proficiency in the weaker language (Xu et al., 2024). A separate study of whole-brain cortical thickness (CT) in the same early and late bilinguals (as in Xu et al., 2024) found that thinner cortex in left and right superior frontal regions was related to greater proficiency in the stronger language (Nguyen et al., 2023). They also reported relationships between thicker cortex in left and right superior frontal regions and left and right parietal regions—including the left and right inferior parietal cortices —and greater proficiency in the weaker language. Additionally, thinner cortex in the pericalcarine cortex was related to greater proficiency in the weaker language (Nguyen et al., 2023). Conversely, Claussenius-Kalman et al. (2021) found no relationships between CT and language proficiency (expressive picture vocabulary) for either of the two languages in a group that included early and late bilinguals. To our knowledge, there is only one study conducted in exclusively *early* bilinguals and it found a positive relationship between CT and language proficiency (receptive picture vocabulary) in left frontal, left and right parietal, and right temporal regions (Vaughn et al., 2021), which was similar to findings reported by Nguyen et al. (2023) in the weaker language. Taken together, several studies found relationships between brain structure and language proficiency, often in left frontal and left and right parietal regions, and these were generally interpreted as experience-dependent effects on brain structure resulting from the presence of another language.

Turning to those studies investigating language balance, Archila-Suerte and colleagues (2018) used a between-group comparison of balanced and unbalanced early bilinguals (balance measured by the difference in vocabulary tests in each of the two languages). They found that balanced early bilinguals had more GMV than the unbalanced bilinguals in ROIs in bilateral putamen, and less CT in ROIs in left middle frontal, inferior frontal, and superior temporal regions (but no differences in surface area). Since the balanced and unbalanced early bilingual groups shared a similar language history and socio-economic environment, the authors suggested the GMV results could be attributed to innate language-learning “aptitude” in the more balanced group (Archila-Suerte et al., 2018), which allowed them to reach equal, or relatively equal, proficiency in both languages. However, the two groups not only differed in their language balance score, but also in language proficiency, raising the question as to whether (i) the structural differences can unambiguously be attributed to language balance to begin with, and (ii) aptitude plays a role. Another study on early bilinguals examined the relationship between GMV and a composite “bilingualism score” based on balance plus proficiency (measured by overall perceived level of comprehension, reading, fluency, pronunciation, writing, and comfort and language use) (Marin-Marin et al., 2022). Collapsing findings across hemispheres, they showed positive correlations between the bilingualism score and GMV in subcortical and cerebellar regions, in addition to a non-linear relationship between bilingualism score and GMV in one inferior frontal region. However, the blending of balance and proficiency (along with other dimensions of bilingualism) into one score obscures the independent effects of either dimension, which presents a challenge for the interpretation similar as in Archila-Suerte et al., (2018).

Claussenius-Kalman et al. (2021) generated a “bilingual proficiency” score, that summed language proficiency for the two languages (expressive vocabulary) and boosted the score based on balance in a mix of early and late bilinguals. Using ROIs, they found thinner cortex for more balanced late bilingual adults in left and right frontal areas. To address proficiency, they tested for associations between CT and language proficiency in these same ROIs and found no significant results, concluding that the findings for CT and balance proficiency could not be attributed to language proficiency. The authors interpreted the findings in left anterior prefrontal cortex and right anterior cingulate cortex in terms of language and cognitive control (required for utilizing and switching between two languages) resulting from language-learning experiences (rather than aptitude).

Taken together, there is some evidence that language proficiency and language balance have a neuroanatomical signature in bilinguals in bilateral frontal and subcortical structures. It is important to note, though, that the results are based on mostly late bilinguals and most are based on a single brain metric. Additionally, almost all studies employed an ROI approach which restricts the inquiry to brain regions that align with a framework proposing that managing two languages results in anatomical growth in brain structures that support language and/or cognitive control. The goal of the current study was to further extend this work by considering relationships between brain structure and both language proficiency and language balance, while also directly controlling the latter with the former. Instead of late bilinguals, we studied *early* cultural bilinguals of Spanish and English who acquired their languages as a part of their environment prior to age six years, thereby reducing heterogeneity of the group. Despite their good language skills, these early bilinguals nevertheless exhibit a range in individual variability of their language proficiency scores and in their language balance scores. Unlike most prior studies, we measured both GMV and CT and did not constrain the voxel/vertex-wise analyses to any regions a priori. Based on the literature, we expected to find relationships between brain structure and language proficiency, and brain structure and language balance, in the left and right frontal and inferior parietal regions, as well as subcortical regions. Importantly, once language proficiency is controlled for in the correlations with language balance, not all of these relationships are likely to survive. Together, the results will advance our understanding of how neuroanatomy relates to language proficiency and language balance in early bilinguals.

## 2. Methods

### 2.1 Participants

Forty-six early Spanish-English bilingual children and adults (average age 16.7 ± 6.6 years; see Table 1) were recruited and tested through methods and materials approved by the Georgetown University Institutional Review Board in line with the principles of the Declaration of Helsinki. Fliers and information were distributed at community events, including local bilingual schools, and around the university campus and surrounding community. Written informed consent was obtained from all (for children, parent consent and child assent was obtained) and data from these same participants have been reported previously in a study comparing them to monolinguals (Schug et al., 2022). None had a history of neurological or psychological disorders and all but four were right-handed as measured by the Edinburgh Handedness Test (Oldfield, 1971). All but two of the participants were ethnically Hispanic, and all but one was exposed to Spanish from birth. Exposure to English occurred between birth and six years of age, on average at around age two. Approximately one-third of the bilinguals were simultaneous bilinguals (exposed to both languages since birth). At the time of study, all participants were using both of their languages on a daily basis, with most reporting using English more often than Spanish and no participant was proficient in any additional language.

**Table 1.** Participant demographics of Spanish-English early bilinguals.

| **N = 46** | **Mean ± SD** | **Range** |
| --- | --- | --- |
| **Sex (M/F)** | 15/31 | - |
| **Age (years)** | 16.7 ± 6.6 | 7.7 — 28.6 |
| **Weaker language proficiency** | 106.7 ± 10.2 | 91 — 136 |
| **Stronger language proficiency** | 121.8 ± 11.1 | 102 — 151 |
| **Language balance** | 15.0 ± 10.6 | 1 — 36 |

*Note:* Counts reported for group size and sex (M/F), means, standard deviation (SD) and range reported for age, weaker language, stronger language, and language balance score (stronger language scores minus weaker language scores).

Analogous, objective measures for language proficiency were obtained in each language using aloud word naming. In Spanish, the *Identificación de letras y palabras* (‘Word-ID’) subtest from the Woodcock-Muñoz *Pruebas de Aprovechamiento III* (Muñoz-Sandoval et al., 2005), and in English, the Letter-Word Identification (‘Word-ID’) subtest of the Woodcock-Johnson III Tests of Achievement were used (Mather & Gregg, 2001; Woodcock, 1987). These tests use single real-word presentation and are standardized and age-normed such that a standard score of 100 represents the average and standard scores from 85 to 115 represent the normal range.

To be included in the study, participants had to have standardized scores equal to or above 85 in both languages. For each participant, the lower of the two scores was labeled the ‘weaker language’ and the higher score was labeled ‘stronger language’. As expected, the average standard language proficiency scores for the group as a whole were within or above the normal range for both the weaker language (*M* = 106.7 ± 10.2; range = 91 – 136) and the stronger language (*M* = 121.8 ± 11.1; range = 102 – 151; see Table 1). The language balance score was calculated for each participant by subtracting their weaker language score from their stronger language score. While all participants performed in the normal to above normal range in both of their languages, the Balance Scores ranged from 1 (very balanced) to 36 (less balanced) standard points; and the average Balance Score was 15.0 (see Table 1). Language balance was negatively correlated with weaker language proficiency (*r* = −0.409, *p* < 0.05) and positively correlated with stronger language proficiency (*r* = 0.516, *p* < 0.05) as would be expected.

Notably, however, the midpoint between the two language proficiency scores was not correlated with the language balance score, indicating that those who are balanced are not necessarily in the higher language proficiency range.

### 2.2 MRI Data Acquisition

A 3T Siemens Trio scanner at the Center for Functional and Molecular Imaging, Georgetown University Medical Center was utilized for image acquisition. High-resolution T1-weighted MR images were obtained using the following parameters: voxel size=1mm×1mm×1mm, TR = 1600 ms, TE = 3.37 ms, flip angle = 15°, field-of-view = 256 mm.

### 2.3 MRI Data Analysis

Blinded to participant demographics, researchers assessed all participants’ raw MRI images for quality. Using a scale of 1 (optimal image) to 5 (severely distorted image), images were rated and no participants were excluded due to poor image quality (images rated above 3).

For GMV, preprocessing of images was performed in SPM12 software using the automated Voxel-Based Morphometry (VBM) technique (Ashburner, J. & Friston, 2000) and methods (Ashburner, J., 2015). The images were segmented into gray matter (GM), white matter (WM), and cerebrospinal fluid (CSF) images and then used to create a study-specific template and spatially normalized to the Montreal Neurological Institute (MNI) stereotaxic space via affine registration of the generated template to the MNI template using DARTEL (Ashburner, 2007). Images were examined to confirm the accuracy of the normalization process and then smoothed with a Gaussian kernel of 10 mm full width at half maximum and the intensity thresholding of the images was set to 0.2.

For CT, image preprocessing was completed using the CAT12 program run through SPM12 software, following the steps outlined in the Computational Anatomy Toolbox (CAT12) Manual (Dahnke et al., 2013). The Images were segmented into separate GM, WM, and CSF images and normalized using the CerebroMatic (COM) Toolbox that allows template formation for samples including both children and adults (Wilke et al., 2017). Following template creation, two quality checks were completed: automatic image quality check through CAT12, for which all images had to score 70% or higher in image quality to be included, and a manual check in which any distorted images were excluded. Images from all participants in the current study passed all quality checks. The images were then smoothed with a Gaussian kernel of 10 mm full width at half maximum.

### 2.4 MRI Correlational analyses

Voxel/vertex-wise statistical analyses of GMV and CT were performed at the level of the whole brain using the SPM12 software. For GMV analyses, correlations were performed for GMV with weaker language proficiency, with stronger language proficiency, and with language balance, all using chronological age and total GMV as covariates of no interest. The correlation between GMV and language balance was repeated after adding weaker language proficiency as a further covariate of no interest. For CT, similar correlations were performed between CT and weaker language proficiency, stronger language proficiency, and language balance, the latter conducted twice, first without any covariates of no interest (age is taken into account statistically in the CT preprocessing pipeline) and then with weaker language proficiency score as a covariate of interest. All correlational analyses were performed in SPM12 using MATLAB version 2021b (voxel-wise height threshold of p < 0.005, FDR; cluster-level extent threshold of p < 0.05).

## 3. Results

### 3.1 GMV and language proficiency

As shown in Figure 1 (and Table 2), there was a positive correlation between GMV and language proficiency in the weaker language in the right angular gyrus (BA 39) extending to the right superior temporal gyrus (BA 22), and no negative correlations. There were no correlation results for GMV with stronger language proficiency score.

**Figure 1.**
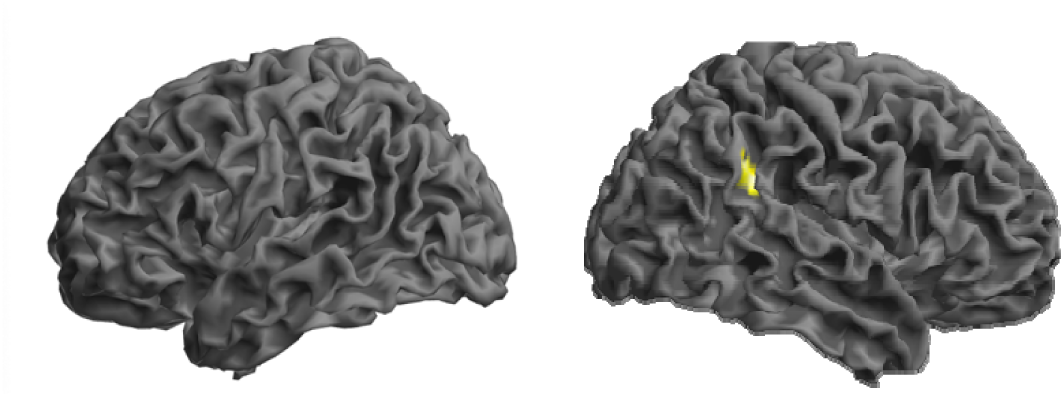
Correlations between GMV and language proficiency scores. More GMV was related to stronger language proficiency in the weaker language. Voxel-wise height threshold p < .005. FDR cluster-level extent threshold p < .05. Coordinates for significant clusters are in Table 2.

**Table 2.**
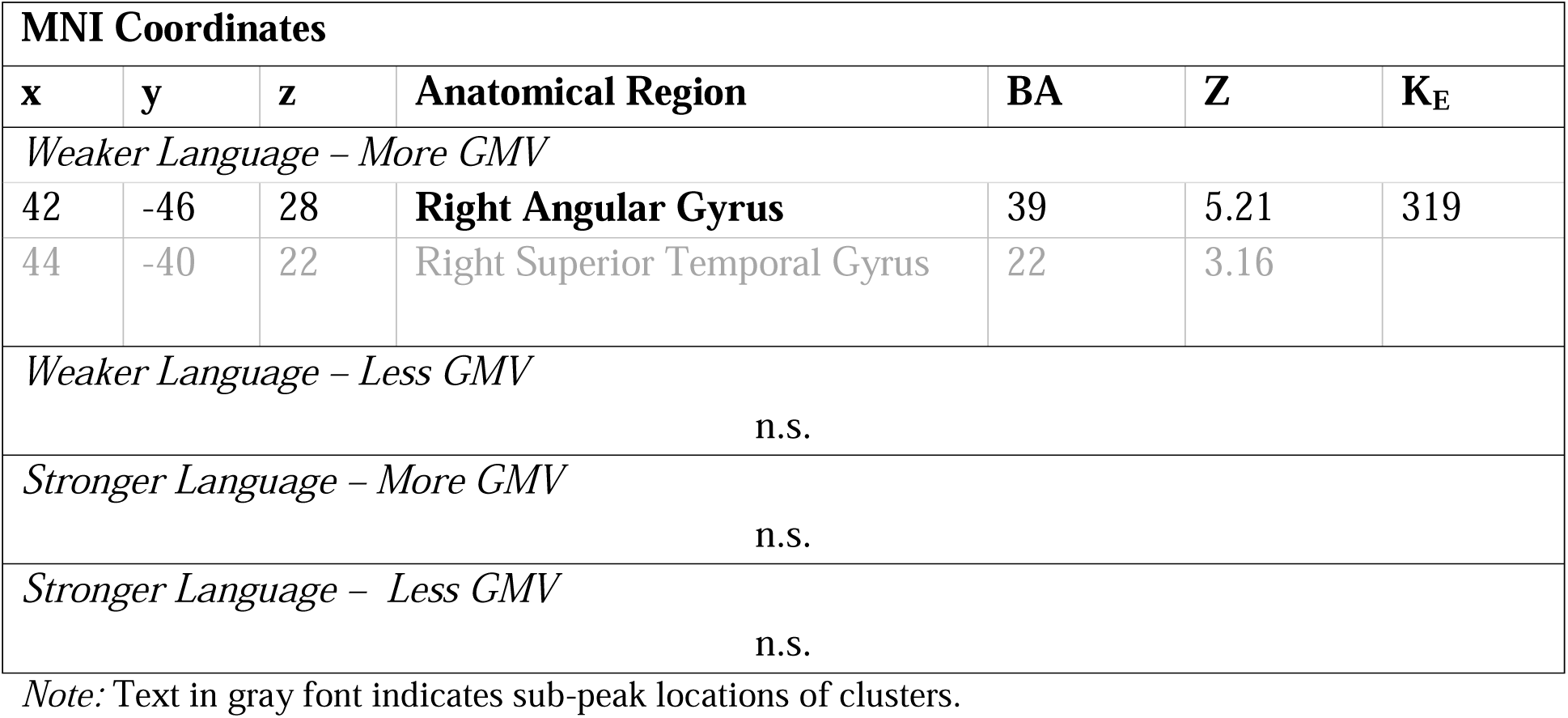
MNI coordinates for maxima of GMV clusters exhibiting a relationship with language proficiency in the weaker and stronger languages.

| MNI Coordinates |  |  |  |  |  |  |
| --- | --- | --- | --- | --- | --- | --- |
| x | y | z | Anatomical Region | BA | Z | K<sub>E</sub> |
| *Weaker Language – More GMV* |  |  |  |  |  |  |
| 42 | -46 | 28 | **Right Angular Gyrus** | 39 | 5.21 | 319 |
| 44 | -40 | 22 | Right Superior Temporal Gyrus | 22 | 3.16 |  |
| *Weaker Language – Less GMV* |  |  |  |  |  |  |
| n.s. |  |  |  |  |  |  |
| *Stronger Language – More GMV* |  |  |  |  |  |  |
| n.s. |  |  |  |  |  |  |
| *Stronger Language – Less GMV* |  |  |  |  |  |  |
| n.s. |  |  |  |  |  |  |

*Note:* Text in gray font indicates sub-peak locations of clusters.

### 3.2 GMV and language balance

Negative correlations between GMV and language balance were found in the right angular gyrus (BA 39), indicating that more balanced bilinguals had *more* GMV in this region (a lower score for language balance signifies more balanced bilingualism). There was a positive correlation between GMV and language balance score in three clusters, indicating more balanced bilinguals had less GMV here: left postcentral gyrus (BA 1) extending to left Rolandic operculum (BA 6); right superior occipital gyrus (BA 18; 19) extending to the right cuneus (BA 18); and, right cerebellar lobule IX extending to the left cerebellar lobule IX (see Figure 2 and Table 3). No significant clusters, however, emerged after controlling for language proficiency (i.e., adding weaker language proficiency score as a covariate of no interest).

**Figure 2.**
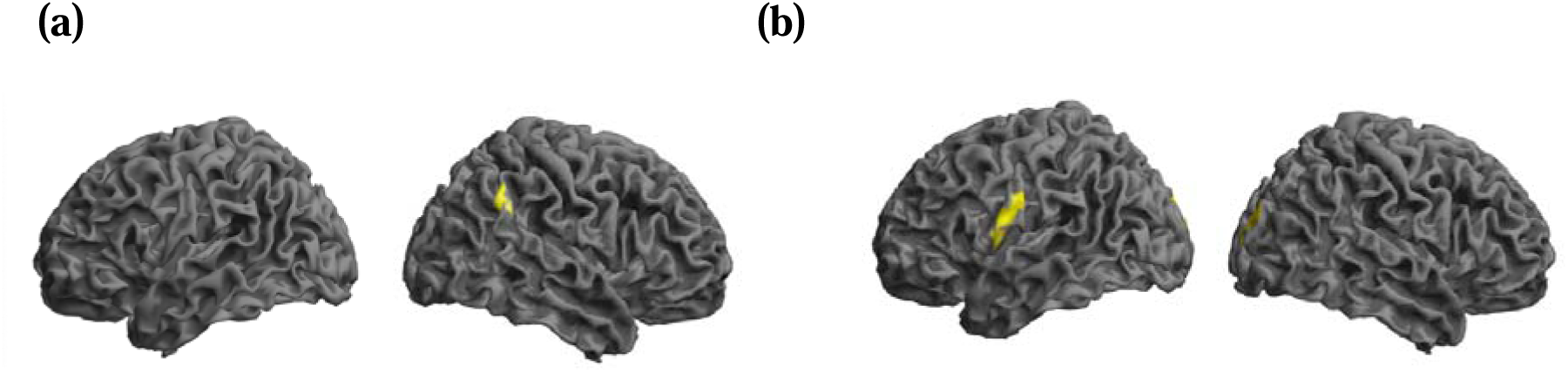
Correlations between GMV and language balance scores without controlling for language proficiency. More balanced bilingualism was associated with areas of (a) more GMV and (b) less GMV Clusters in the cerebellum not visualized in figure. Voxel-wise height threshold p < .005. FDR cluster-level extent threshold p < .05. Coordinates for significant clusters are in Table 3.

**Table 3.** MNI coordinates for maxima of GMV clusters where there is a relationship with language balance, both without and with a control for weaker language proficiency.

| **MNI Coordinates** |  |  |  |  |  |  |
| --- | --- | --- | --- | --- | --- | --- |
| **x** | **y** | **z** | **Anatomical Region** | **BA** | **Z** | **K<sub>E</sub>** |
| *Language balance (without control for proficiency) – More GMV* |  |  |  |  |  |  |
| **50** | **-50** | **36** | **Right Angular Gyrus** | **39** | **3.58** | **328** |
| 62 | -57 | 34 | Right Angular Gyrus | 39 | 2.85 |  |
| *Language balance (without control for proficiency) – Less GMV* |  |  |  |  |  |  |
| **-64** | **-16** | **30** | **Left Postcentral Gyrus** | **1** | **4.16** | **762** |
| -66 | -10 | 24 | Left Postcentral Gyrus | 1 | 3.82 |  |
| -54 | -3 | 12 | Left Rolandic Opercularis | 6 | 3.77 |  |
| **4** | **-63** | **-60** | **Right Cerebellum Lobule IX** | **-** | **3.58** | **425** |
| -2 | -54 | -62 | Left Cerebellum Lobule IX | - | 3.49 |  |
| 6 | -52 | -64 | Right Cerebellum Lobule IX | - | 3.19 |  |
| **14** | **-102** | **18** | **Right Superior Occipital Gyrus** | **18** | **3.51** | **429** |
| 28 | -94 | 26 | Right Superior Occipital Gyrus | 19 | 3.38 |  |
| 18 | -93 | 9 | Right Cuneus | 18 | 3.38 |  |
| *Language balance (controlling for proficiency) – More GMV* |  |  |  |  |  |  |
| n.s |  |  |  |  |  |  |
| *Language balance (controlling for proficiency) – Less GMV* |  |  |  |  |  |  |
| n.s |  |  |  |  |  |  |

*Note:* Text in bold font denotes the peak of the cluster and italic font indicates any sub-peak locations within each cluster.

### 3.3 CT and language proficiency

No significant clusters were revealed in correlations between CT and weaker language proficiency and in correlations between CT and stronger language proficiency.

#### 3.3.4 CT and language balance

Correlations between CT and language balance revealed no significant clusters, without and with weaker language proficiency as a covariate of interest.

## 4. Discussion

Several prior studies in bilinguals have tested for brain structure-behavior relationships for language proficiency (Mechelli et al., 2004; Grogan et al., 2012; Xu et al., 2024; Nguyen et al., 2023; Claussenius-Kalman et al., 2021;Vaughn et al., 2021) or language balance (Archila-Suerte et al., 2018; Marin-Marin et al., 2022; Claussenius-Kalman et al., 2021), but rarely together (Claussenius-Kalman et al., 2021), even though the interpretation of the latter depends on the former. The aim of the current study was to build on this disparate literature by jointly testing for relationships between brain structure and both language proficiency, and language balance. Further, this study employed a whole-brain, voxel/vertex-wise approach, whereas all prior studies noted above except one (Nguyen et al., 2023) spatially constrained their brain analyses, primarily through ROIs. It employed two brain structure metrics (GMV, CT), given the importance of using multiple metrics in the same study (Claussenius Kalman et al., 2020). Also, unlike prior studies, this investigation focused on early cultural bilinguals, which provides for a more homogenous group of bilinguals who learn both of their languages through their family environment rather than classroom-based instructions.

Our results revealed a positive relationship between GMV and language proficiency in the right angular gyrus extending into superior temporal gyrus, and that same right angular gyrus region exhibited more GMV in those with greater language balance. There were also regions with the opposite pattern, revealing less GMV for more balanced bilinguals. However, once language proficiency was controlled for in these correlation between GMV and language balance, these findings disappeared. Turning to CT, there were no significant relationships between CT and language proficiency nor CT and language balance. These results provide insight into the role of individual differences in language performance within early bilinguals. Notably, they reveal that language proficiency is related to brain regions outside those tied to language, suggesting a more domain-general mechanism for stronger proficiency; and that once proficiency is accounted for, there is no evidence that brain anatomy is related to proficiency discrepancies between the two languages. They also demonstrate that the results are dependent on the neuroanatomical metric used.

### 4.1 Early bilinguals’ proficiency is positively correlated with GMV in the right angular gyrus

We found that greater GMV in the right angular gyrus/superior temporal gyrus was associated with higher proficiency in the weaker language. For context with the prior literature (primarily conducted in late bilinguals), only one (Grogan et al., 2012) of the three GMD/GMV studies (Mechelli et al., 2004; Grogan et al., 2012; Xu et al., 2024;) included a right inferior parietal region (the supramarginal gyrus) as an ROI, given that the focus for most investigations was on left hemisphere inferior frontal and/or inferior parietal regions. However, the correlation analysis between GMD and GMV with second language proficiency in right supramarginal gyri yielded null findings (Grogan et al. 2012). They interpreted their findings in left inferior frontal cortex in terms of neuroplasticity related to language skills, specifically the acquisition of a larger vocabulary (Green et al., 2007). While GMV in right parietal cortex is associated with naming of words in prior work (Johns et al., 2018), the right angular gyrus is not commonly associated with language, but rather with attentional and control processes (Seghier, 2013) and sensemaking. This is defined as the neural process of giving meaning to external sensory information and involves defining the current context, allocating attentional resources, extracting relevant information, supporting social cognition and converging toward plausible accounts of events and contexts, and may be related to bilingual language contexts and use (Price & Mechelli, 2005; Seghier, 2023; Wang et al., 2020). This association suggests that for early bilinguals (rather than late bilinguals who likely learn a second language through more explicit language learning efforts), higher proficiency in their weaker language may be less related to processes such as vocabulary learning in language regions, and more dependent on domain-general processes in the right angular gyrus.

Prior research utilizing the same bilingual sample as in the current study found differences in GMV between this group and a monolingual, such that the bilinguals had relatively more GMV in a region extending from the right postcentral gyrus to the right supramarginal gyrus of the inferior parietal cortex (Schug et al., 2022). This finding (BA 40; x= 48 y= −33 z= 46) is near the right angular gyrus finding revealed in this investigation to be positively correlated with language proficiency (BA 39; x= 42, y= −46, z= 28). Thus, it is plausible that the IPC is involved in the structural reorganization associated with a dual-language experience in ways that positively relate to proficiency of the weaker language.

While the right angular gyrus finding has no precedence in the literature, this may be due to the common practice of ROI analyses. Mechelli et al. (2004), identified positive associations between brain structure and language proficiency in left angular gyrus, which Grogan et al. (2012) were not able to replicate, but instead reported a finding in left frontal lobe, while Xu et al., 2024 focused their investigation on subcortical regions and found GMV there to be related to language proficiency. Taken together, prior GMD/GMV studies show no convergence due to lack of whole-brain studies.

### 4.2 There is a relationship between GMV and language balance, but not after accounting for language proficiency

Correlations revealed relationships between more GMV in the right angular gyrus for more balanced early bilinguals, while balanced early bilinguals also had less GMV in the left postcentral gyrus/Rolandic operculum, right superior occipital gyrus/cuneus and bilateral cerebellar lobule IX. The right angular gyrus finding coincided with the result from the correlation between GMV and language proficiency. Since left postcentral gyrus (Behroozmand et al., 2015) and right cerebellum (Chen et al., 2025; Stoodley et al., 2021) play a role in speech and verbal fluency, it might be tempting to speculate that less GMV in brain regions which support general aspects of fluency and articulation might hold the key to becoming a balanced bilingual, instead of those considered to be more directly involved in language and language control. However, all these findings disappeared once we accounted for language proficiency. As such our results indicate that there is no evidence for a neuroanatomical region that facilitates balanced bilingualism, other than those already implicated in supporting language proficiency.

A prior study on language balance used a between-group comparison and an ROI analysis of the caudate and putamen, regions implicated in inhibition and language switching, and found that “balanced” and “unbalanced” bilingual children differed in GMV of the putamen such that the balanced bilinguals had relatively more GMV in this region (Archila-Suerte et al., 2018). However, their groups also differed in proficiency, leaving open the possibility that proficiency, rather than balance per se, drove these GMV differences. Marin-Marin et al. (2022) also found a positive association between GMV in the putamen and bilingualism score, but here, too, balance was not differentiated from language proficiency. No basal ganglia structures emerged from our whole-brain analysis but given the role of basal ganglia structures in the literature for language proficiency (Xu et al., 2024) and for language balance (Archila-Suerte et al., 2018; Marin-Marin et al., 2022) we conducted a post-hoc ROI analysis of the putamen using the AAL Atlas (Rolls et al., 2020), but there found no significant relationships.

Our results indicate that language balance does not exhibit a distinct neuroanatomical relationship once language proficiency is taken into account, because GMV variability related to language balance is largely shared by language proficiency. We have no evidence that there are brain structures that manifest an influence beyond that related to language proficiency (such as one associated with aptitude, as reported by Archila-Suerte et al., 2018). These results highlight the importance of modeling language proficiency and balance simultaneously when investigating bilingual neuroanatomy.

### 4.3 There is no relationship between CT and language proficiency nor balance

In contrast to the GMV findings, we found no significant relationships between CT and either language proficiency or language balance, regardless of whether proficiency was controlled for. Prior studies, however, have reported such relationships. Nguyen et al. (2023) identified relationships between CT and language proficiency for both languages: thinner cortex in left and right frontal regions was associated with higher language proficiency in the stronger language, while thicker cortex in other left and right frontal and parietal regions, as well as in the pericalcarine cortex, was associated with higher language proficiency in the weaker language.

This finding in the weaker language largely converges with results from Vaughn et al. (2021), using parcellated data in early bilinguals from the ABCD Study dataset, which consist of positive associations between CT in left frontal, left and right parietal, and right temporal regions and language proficiency. Given this agreement, null results for relationships between CT and language proficiency from another study (Claussenius-Kalman et al., 2021) and the present study may therefore be surprising, especially since the present study observed brain-behavioral relationships with GMV. This reinforces the importance of examining multiple metrics when investigating brain structure in bilinguals (Claussenius-Kalman et al., 2021). Notably, even though Claussenius-Kalman and colleagues (2021) did not find a relationship between CT and language proficiency, they did find a relationship between CT and “bilingual proficiency.” Specifically, thinner cortex in ROIs in left and right frontal areas was associated with more balanced bilingualism. Given the few studies conducted on the relationship between CT and language balance in bilinguals, more work is warranted.

### 4.4 Future Studies

Novel aspects of the present study are to test for brain-behavioral correlations simultaneously for both language proficiency and language balance, the use of whole-brain analyses and two, rather than a singular, structural metric, and the focus on early cultural instead of late bilinguals. While it was not our goal to understand these relationships in different types of bilinguals, future studies could conduct a joint investigation into both early as well as late bilinguals, to test if a dual-language experience manifests differently in these two groups in terms of language balance. Also, future studies could benefit from expanding further on the measures used for brain anatomy and include brain function.

## 5. Conclusions

In sum, the present study demonstrates that in early cultural Spanish-English bilinguals, individual differences in language proficiency are related to right angular gyrus GMV, known to serve attentional and control processes rather than language. While individual differences in language balance also are related to GMV in the right angular gyrus, once language proficiency is accounted for, the association with balance disappears, underscoring the importance of examining these language dimensions jointly, rather than attributing brain structural variability to balance alone. There were no results for the measure of cortical thickness, underscoring the need of using multiple measures of brain anatomy. These findings support a graded view of bilingualism in which individual differences in proficiency, even among individuals who acquired both languages early and to a high level of proficiency, are associated with experience-dependent structural variation.

## Acknowledgements

The authors thank the participants and their families for their time and commitment. The authors would like to acknowledge Breana Downey, Melanie Lozano, Nicole Schlosberg, Cambria Revsine, Lynn Flowers, Kelly Mandella, Ryan Mannion, Eileen Napoliello, Ashley Piche for their contribution to the data acquisition and Alison Schug for input on the data analyses.

## Declaration of generative AI use

No generative AI was used in the manuscript preparation process.

